# Early eukaryotic origins and metazoan elaboration of MAPR family proteins

**DOI:** 10.1101/737684

**Authors:** Elisabeth Hehenberger, Michael Eitel, Sofia A.V. Fortunato, David J. Miller, Patrick J. Keeling, Michael A. Cahill

## Abstract

**Background:** The membrane-associated progesterone receptor (MAPR) family consists of heme-binding proteins containing a cytochrome b_5_ (cytb_5_) domain characterized by the presence of a MAPR-specific interhelical insert region (MIHIR) between helices 3 and 4 of the canonical cytb5-domain fold. Animals possess three MAPR families (PGRMC-like, Neuferricin and Neudesin).

**Results:** Here we show that all animal MAPR families were already present in the common ancestor of the Opisthokonta (comprising animals and fungi as well as related protistan taxa). All three MAPR genes acquired extensions C-terminal to the cytb_5_ domain, either before or with the evolution of animals. The archetypical MAPR protein, progesterone receptor membrane component 1 (PGRMC1), contains phosphorylated tyrosines Y139 and Y180. The combination of Y139/Y180 appeared in the common ancestor of Cnidaria and bilaterally symmetrical animals, along with an early embryological organizer and synapsed neurons, and is strongly conserved in all bilateral animals. A predicted protein interaction motif in the PGRMC1 MIHIR is potentially regulated by Y139 phosphorylation. A multilayered model of animal MAPR function acquisition includes some pre-metazoan functions (e.g., heme binding and cytochrome P450 interactions) and some acquired animal-specific functions that involve regulation of strongly conserved protein interaction motifs acquired by early-branching animals.

**Conclusions:** This study provides a conceptual framework for future studies, against which PGRMC1’s multiple functions can perhaps be stratified and functionally dissected. In accompanying papers we show that mutational perturbation of PGRMC1 phosphorylation status of the Y180 motif is associated with dramatic changes cell pasticity assayed by protein abundances, cell morphology, mitochondrial function, genomic stability, and epigenetic status, with pathways analysis associating Y180 mutation with processes related to organizer function. These combined works reveal previously unrecognized involvement of PGRMC1 in foundational animal processes of great relevance to disease.

## BACKGROUND

Progesterone receptor membrane component 1 (PGRMC1) is the archetypal member of the membrane associated progesterone receptor (MAPR) family [1, 2]. Vertebrates including humans encode four MAPR proteins. PGRMC1 and the closely related PGRMC2 arose by gene duplication of an original ‘PGRMC’ gene during vertebrate evolution. We refer here to PGRMC1 and/or PGRMC2 for proteins from vertebrates that have inherited this gene duplication [3, 4], or otherwise to PGRMC. Other vertebrate MAPR proteins are neuron-derived neurotrophic factor, commonly known as neudesin (NENF), and neuferricin (NEUFC) [3–6].

PGRMC1 has a long list of seemingly disparate functions, ranging from involvement in steroid and heme synthesis, membrane trafficking, progesterone response in fertility and other situations, and conferral of progesterone-dependent anti-apoptosis [7]. PGRMC1 guides embroyonic axonal growth in nematodes and mammals [8, 9] (i.e. perhaps in all bilaterally symmetrical animals), is expressed in a variety of neurons of the central nervous system (CNS) [10–13], and is found in synpases where it affects membrane trafficking [14]. NENF is present in CNS regions of embryonic neural differentiation [15, 16]. In the CNS its expression pattern suggests an exclusive role in neurons, while *in vitro* NENF exhibits strong neurotrophic activity as a secreted protein [16]. Like all vertebrate MAPR proteins, NEUFC is implicated with cytochrome P450 reactions, steroidogenesis, and neurobiology [5, 17–19].

All MAPR proteins contain an insertion of an oligopeptide sequence between helices 3 and 4 of the canonical cytochrome b_5_ (cytb_5_) domain fold (as defined by human cytb_5_). In place of a short loop at this position in classical cytb_5_ domain proteins, MAPR proteins contain a MAPR interhelical insertion region (MIHIR) of variable length [1, 4]. The MIHIR of PGRMC1 and PGRMC2, but not of NENF or NEUFC, contains a tyrosine (PGRMC1 Y139) that is strongly conserved in later-branching animals [4].

Interest in PGRMC1 phosphorylation was sparked when it was found to exist in different phosphorylated forms in breast cancers that were positive or negative for estrogen receptor expression [20]. Bioinformatics revealed the presence of two Src homology 2 (SH2) and one Src homology 3 (SH3) domain target motifs (short peptide sequences that would bind to a much larger SH2 or SH3 protein domain, respectively: hereafter SH3 and SH2 motifs), being a proline-rich SH3 motif centered on PGRMC1 P63, and SH2 motifs centred on Y139 and Y180 [2, 21]. Notably, the P63 SH3 motif and the Y180 SH2 motif were flanked by consensus casein kinase 2 (CK2) sites with phosphoacceptors at S57 and S181. CK2 is constitutively active in many cells, contributing to the order of 20% of the human phospho-proteome [22, 23]. This suggested a model where CK2 phosphorylation could sterically prevent interactions of kinases or other interaction partners with the motifs at P63 and Y180, thereby negatively regulating PGRMC1 function [2, 20]. Mutation of both of the CK2 sites, but not each individually, rendered MCF-7 breast cancer cells resistant to peroxide-induced cell death [20]. However, a recent knockout of CK2 activity in C2C12 mouse muscle cells revealed marginally higher phosphorylation of S181 (and Y180), clearly showing that a kinase other than CK2 can target S181 [24].

Phosphorylation of both residues at CK2 consensus sites and of the Y139 and Y180 SH2 motifs, as well as a variety of other residues, has been observed from high throughput proteomics studies [25]. Phylogenetic analysis revealed that PGRMC1 acquired signalling and phosphorylation motifs during animal evolution: e.g., the PGRMC1 SH3 motif is absent from PGRMC2, and was gained by terrestrial tetrapods. The adjacent S57 phosphorylation site appeared during primate evolution [4]. It was previously incorrectly concluded that the ancestral metazoan appears to have possessed cognates of both the PGRMC1 Y139 and Y180 SH2 target motifs [4]. As we demonstrate here, this incorrect conclusion was due to mis-assignment of several early-branching metazoan MAPR proteins to the PGRMC family.

A CK2 consensus site adjacent to Y180 arose in the common ancestor of Bilateria [4], reflecting an embryological state before vertebrate body plan is determined. This is of particular interest to early animal evolution and embryology because, 1) ligands and receptors of the Wnt pathway evolved in early animals [26, 27], and PGRMC1 regulates this pathway in mammalian pluripotent stem cells [28]; 2) Both nematode MAPR proteins are expressed through early nematode embryogenesis from the oocyte stage until the induction of germline segregation and neural differentiation [29]; 3) PGRMC1 is implicated in essential progesterone (P4)/progestin responses in male [30–32] and female [21, 33, 34] germline and reproductive cells; 4) PGRMC-like proteins direct ventral embroyonic neural axon migration conserved between nematodes and mammals [8, 9]; 5) PGRMC1 Y180 phosphorylation was observed only in synaptic fractions of mouse neurons [35]; and 6) PGRMC1 is involved in synaptic membrane trafficking [14], implicating a role of PGRMC1 phosphorylation in the synaptic signaling that serves a key organismic coordination role in all animals with a nervous system.

It remains unclear which function(s) may be regulated by PGRMC1 phosphorylation. PGRMC1 is a multifunctional protein [2, 7]. Heme-binding and cytochrome P450 (cyP450) modulation are properties attested from protist MAPR proteins [6, 36–38]. We reasoned that the plethora of PGRMC1 functions should be separable into ancient eukaryotic roles (such as cyP450 interaction) and newly acquired metazoan roles, such as hypothesized tyrosine phosphorylation-mediated signaling in animals. In the present study we examine the nature of MAPR diversity in early-branching animals as well as in unicellular lineages that represent the closest relatives of animals, with particular interest in the origins of PGRMC1 functional SH2 motifs in opisthokonts.

The Opisthokonta are a eukaryotic supergroup forming two lineages, the Holozoa and the Holomycota [39]. While the Holomycota include fungi and their relatives (such as nucleariids), the Holozoa consists of animals together with their closely related unicellular sister lineages (choanoflagellates, filastereans, pluriformeans and ichthyosporeans) (Figure 1A). Tyrosine kinases and SH2 domains (which bind to phosphorylated tyrosine residues) evolved in those Holozoan unicellular animal relatives, as well as many other proteins normally associated with animals [27, 40–42], including new transcription factors, cell surface adhesion molecules, transposons, and extracellular matrix [43].

**Figure 1.**
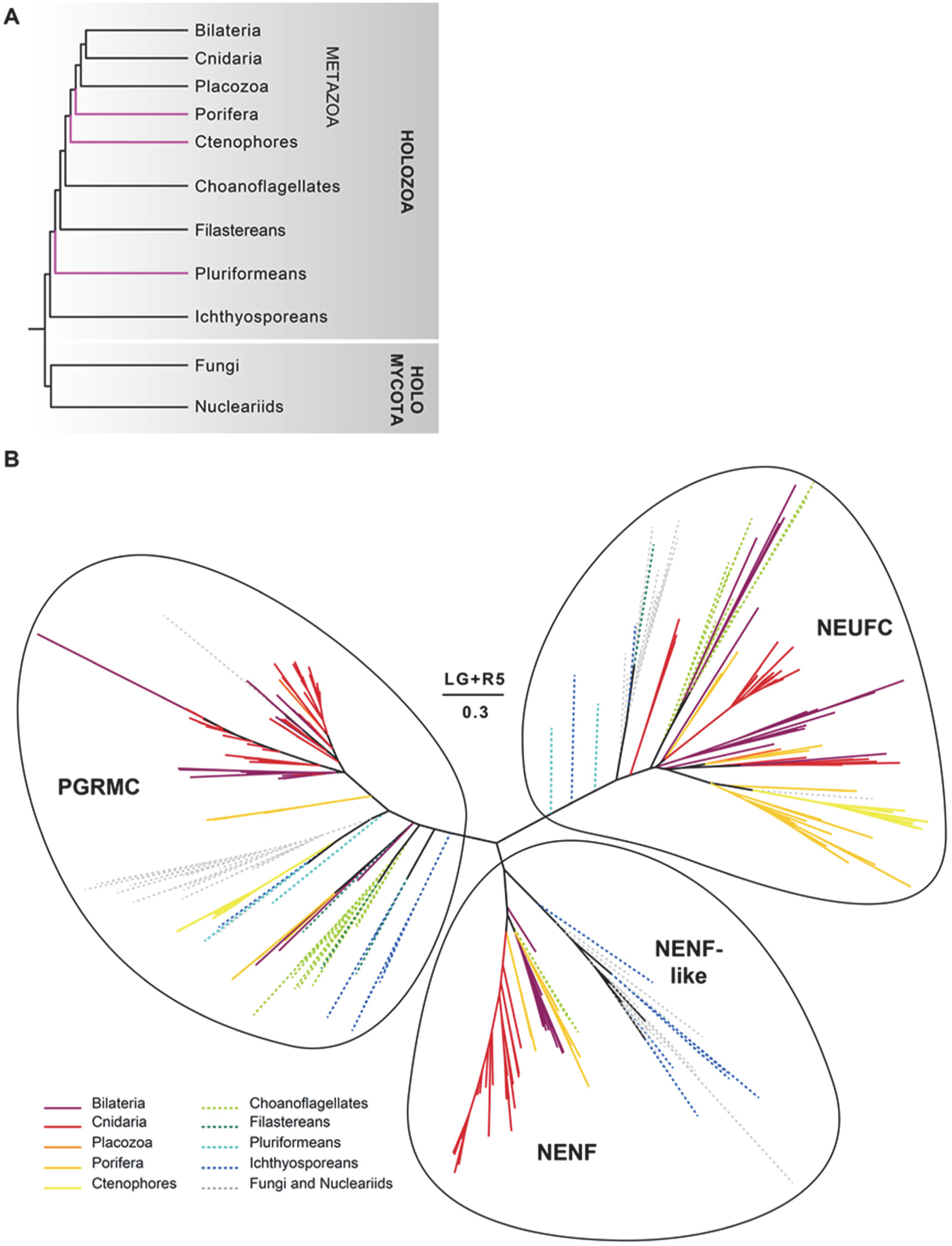
Phylogenetic reconstruction of MAPR proteins in opisthokonts. (A) Schematic tree of opisthokont lineages analyzed in this work, with contentious branches colored in magenta. The branch topology of Ctenophores and Porifera divergence is subject to strong ongoing debate, with both Porifera [88, 105] and Ctenophores [106] argued as forming sister groups to all other animals. Similarly, the Pluriformea (*Corallochytrium*) were found to branch together with Ichthyosporea as sister to all other holozoans in alternative tree reconstructions [107]. (B) Phylogeny of 3 types of MAPR proteins in opisthokonts: progesterone receptor membrane component (PGRMC), neudesin (NENF and NENF-like) and neuferricin (NEUFC). Solid lines represent metazoan lineages, dashed lines represent non-metazoan lineages. Different lineages are indicated by color in the key. The scale bar and the number beneath it indicate the estimated number of substitutions per site, above the scale bar the model for tree reconstruction is indicated. For bootstrap support see Supplemental Figure 1, for a phylogeny containing also taxon information, see figshare repository doi: 10.6084/m9.figshare.9162164.

A previous pilot study [4] of the phylogenetic distribution of animal PGRMC1 suffered from taxonomic bias against early-diverging animals and their unicellular relatives due to insufficient taxon sampling, especially in critical transitions of early animal evolution, and from poor discrimination between MAPR family members in early-branching organisms. The guiding motivation of the present study was to identify the members of the MAPR protein families present in early-branching animals and their protistan relatives, aiming to understand the changes in MAPR proteins, evolutionarily conserved regions of importance, and particularly to better define the acquisition of PGRMC1 functions.

## RESULTS

### The origin of animal MAPR protein families predates Opisthokonta

The relationships of the major opisthokont groups investigated in our analysis are depicted schematically in Figure 1A based upon published phylogenies [44, 45], acknowledging that the tree of Opisthokonta is not resolved (see the legend of Figure 1A for details). Since metazoan evolution has been associated with the evolution of many new membrane proteins relative to intracellular enzymes [46], we were interested in the evolution of individual MAPR protein families, particularly in early-branching animals. We obtained diverse opisthokont MAPR proteins across a wide selection of taxa and investigated their phylogenetic relationships. The unrooted tree topology resulting from our MAPR analysis indicated the presence of three well-supported major branches of opisthokont MAPR proteins, representing PGRMC, NENF and NEUFC proteins [6], with members of each family also present in fungi (Figure 1B and Figure S1). Not all species sampled possessed examples of each gene family (e.g., both yeasts *Saccharomyces cerevisiae* and *Schizosaccharomyces pombe* contain only one MAPR protein: Dap1, a PGRMC family protein). However, this result clearly reveals that the three MAPR families were already distinct in the common ancestor of opisthokonts. Below we describe in detail the characteristics and the evolution of each MAPR protein family.

The consensus Logo plots of the MAPR domains for each individual family from Figure 1B, as well as for all MAPR sequences from all families combined are shown in Figure 2. Human PGRMC1 sequence is shown above the plots for orientation. The location of the MAPR domain in each of the human MAPR family proteins is schematized in Figure 3. Changes to each protein family, particularly along the transition from unicellular to multicellular holozoans as well as non-bilaterian animals to bilatieria, are discussed below. These include changes to the C-termini of each protein family (Figure 3).

**Figure 2.**
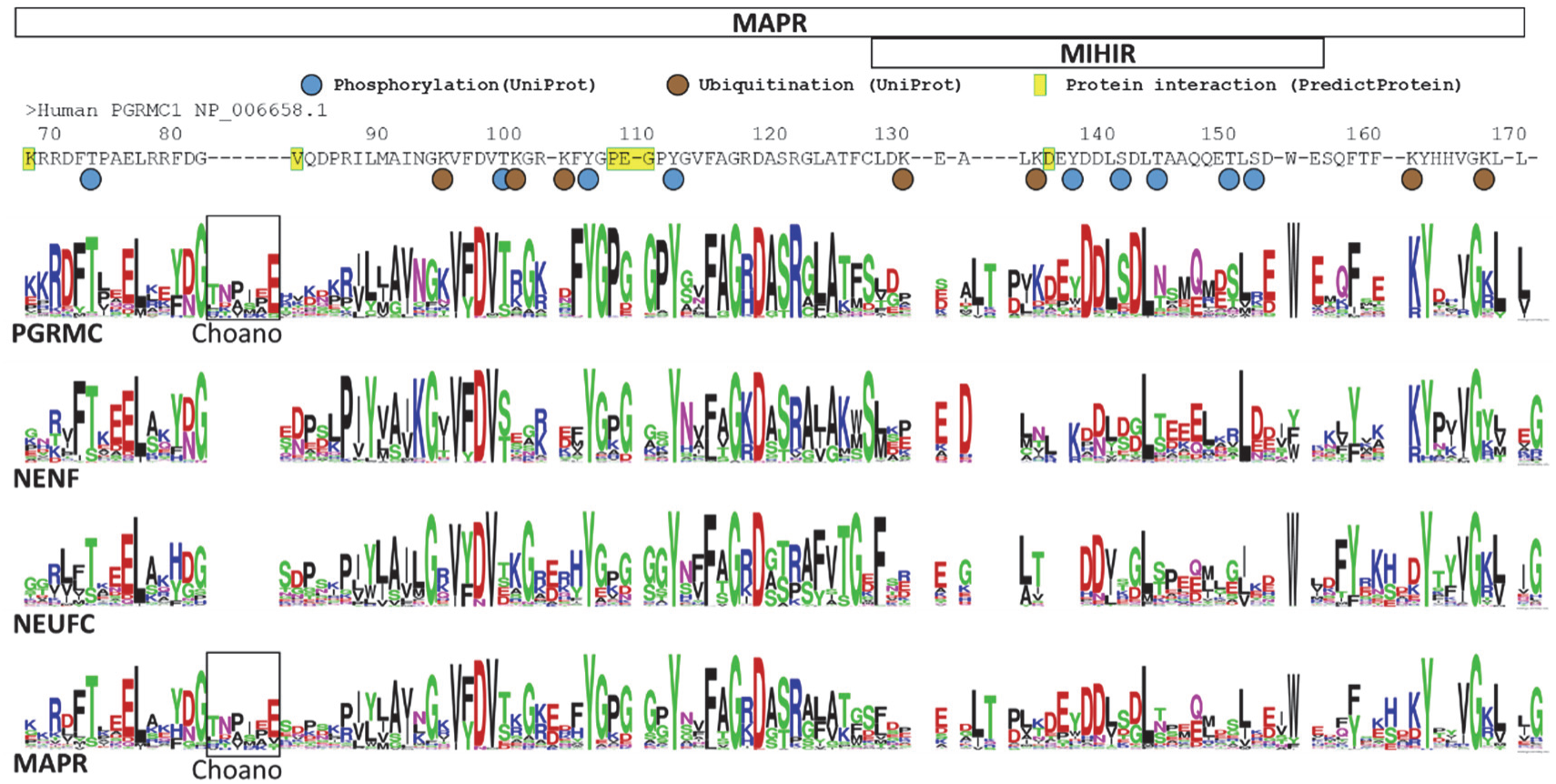
Consensus MAPR and subfamily Logo plots. Logo plots are presented for all members of the PGRMC, NENF, and NEUFC families from Figure 1. The consensus plot of all MAPR sequences in the lowest row highlights the overall MAPR domain sequence identity. Apomorphic sequence insertions from some individual sequences were deleted to facilitate presentation. The box represents an apomorphic insertion in choanoflagellates that is absent from PGRMC proteins of other species. The MAPR domain of human PGRMC1 is presented above the Logo plots for reference. Documented sites of phosphorylation, ubiquitination (UniProt) and predicted sites of interaction (ProteinPredict) are indicated for PGRMC1.

**Figure 3.**
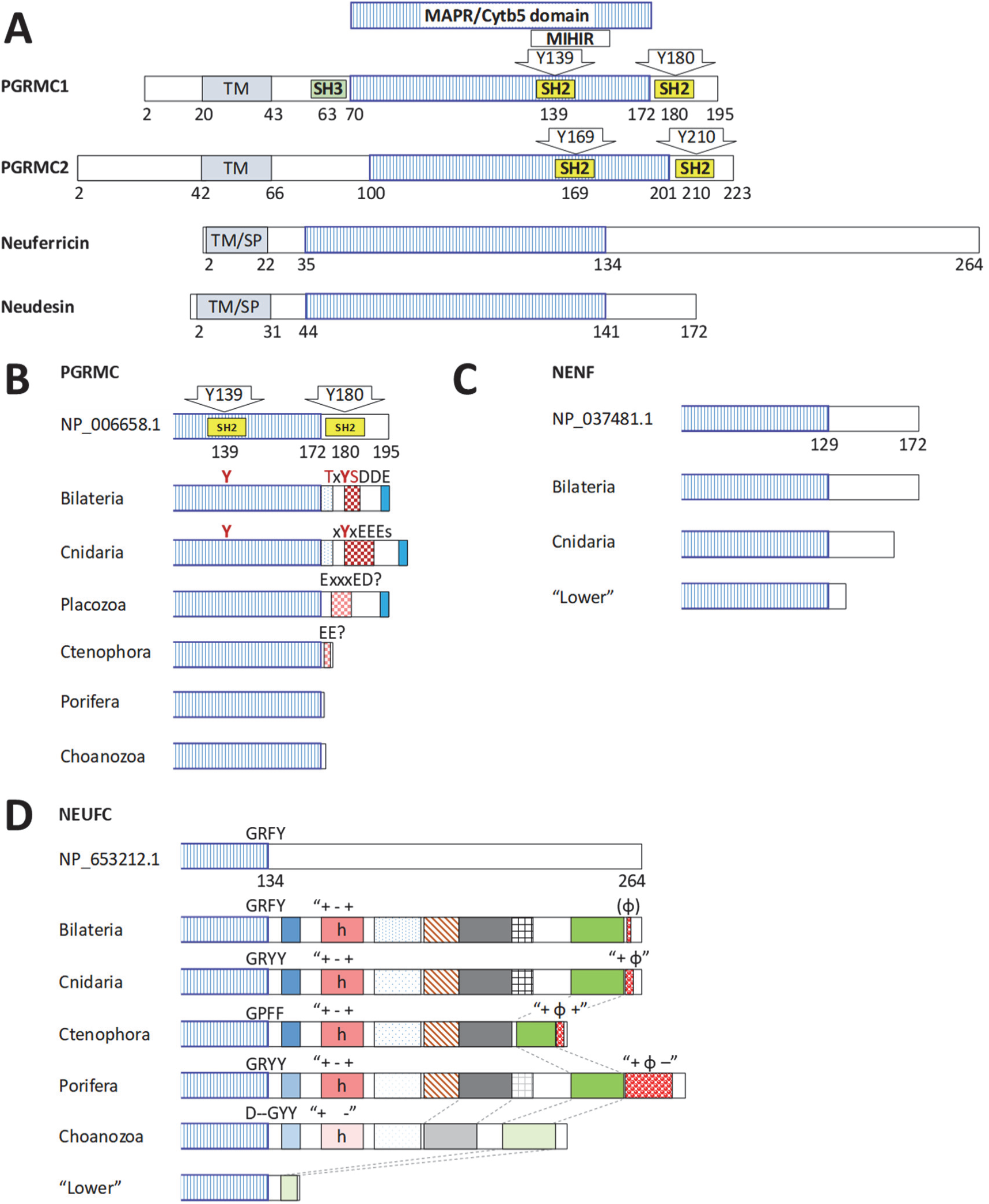
Evolution of MAPR C-termini in the evolution of animals. (A) Schematic depiction of the four human MAPR genes. Numbering refers to human proteins with accession numbers provided in subsequent panels. TM: transmebrane peptide of PGRMC1/2; TM/SP: Transmembrane/signal peptide of Neuferricin and Neudesin [3]. (B) Schematic depiction of the evolution of the PGRMC C-terminus in the evolution from Choanozoa to Bilateria. Human PGRMC1 from A is at the top for orientation. The cognate positions of Y139 and Y180 are shown to have appeared in Cnidaria but are absent from earlier-diverging animals. Boxed regions show regions of amino acid similarity without identifiable known domains. These are not implied to possess specific functions. The PGRMC1 Y180 motif consisting of T178, Y180, S181 and adjacent negative D/E region appears to have evolved in a stepwise pattern. Logo plots and further details of the schematic diagrams can be found in Figure S2. (C) The C-terminus of NENF proteins expanded during the evolution of Cnidaria and Bilateria. Logo plots and further details of the schematic diagrams can be found in Figure S4. “Lower” refers to earlier-branching groups. (D) The C-terminus of NEUFC was expanded in the evolution from earler-diverging opisthokonts to Holozoa (Choanozoa and Metazoa). Variously shaded boxes represent regions of presumed amino acid similarity by descent. No function is ascribed to any particular region. Consensus changes to the GFRY motif at the C-terminus of the human Neuferricin (NP_653212.1) are shown for each group. The boxes labelled h with “+ −“ above represents regions of positive and negative charge that is predicted to be surface-exposed. Another region contains positively (+) or negatively (-) charged and/or aliphatic (ϕ) residues (see Figure S7B-E). Logo plots and further details of the schematic diagrams can be found in Figure S8. “Lower” refers to earlier-branching groups.

### The PGRMC family

Notably, PGRMC proteins feature prominent F106, P109 and P112 (with reference to human PGRMC1) around the heme-binding pocket of the MAPR domain (Figure 2), suggesting unique ligand properties of this family. Choanoflagellates, the closest unicellular sister group to animals, possess an apomorphic PGRMC-specific insertion between PGRMC1 G83 and V84 (Figure 2).

#### N-terminus

There was no observable systematic change of PGRMC protein size N-terminally to the MAPR domain between early- and late-diverging opisthokonts (not shown). PGRMC1 residues 47-49 encode a putative RGD protein interaction motif which has been present at least since the emergence of placental mammals (Figure 4).

**Figure 4.**
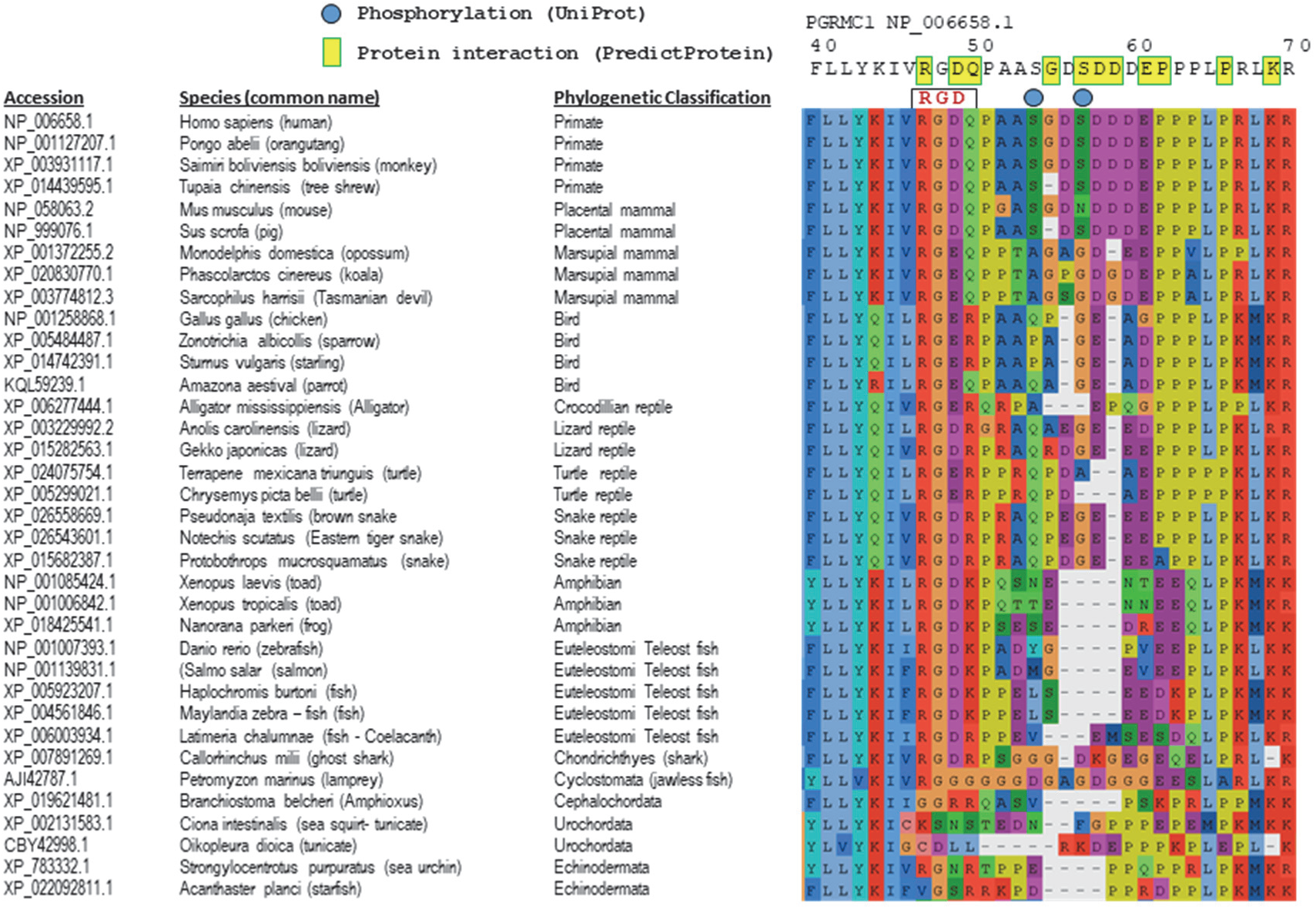
Alignment of PGRMC1 39-74 region of selected chordates. This region spans part of the transmembrane helix (left) to the start of the MAPR domain (right). The indicated metazoan PGRMC sequences were aligned using MAFFT L-INS-i. The graphical presentation of the alignment was made using AliView [108].

#### MAPR domain

In the transition from unicellular holozoans to cnidarians the PGRMC MAPR domain acquired a frequently represented N-terminal KKR consensus corresponding to PGRMC1 69-71. The two placozoan sequences, cnidarians and bilaterians all featured increasing frequency of K96 and K137, which are ubiquitinated in mammals. K96 is also acetylated (Phosphosite [47], action?id=5744). The frequency of tyrosine at the position of SH2 motif Y139 increases markedly in cnidarian and bilaterian animals (Figure 3, Figure S2).

#### C-terminal extension to the MAPR domain

Relative to unicellular Holozoa, Porifera and Ctenophora, a C-terminal extension beyond the PGRMC MAPR domain is present in placozoans, cnidarians and bilateral animals, consistent with closer affiliation of placozoans to later-branching metazoans than to Ctenophora and Porifera [48, 49]. A conspicuous conserved feature corresponds to the terminal 192-195 RKND motif of PGRMC1, which was already recognisable in both placozoan sequences available, and which featured strongly in cnidarian and bilaterian proteins (Figure S2).

The most prominent characteristic of the bilaterian PGRMC C-terminus corresponds to the PGRMC1 TxYSxDDE motif surrounding Y180, where phosphorylation of T and S is postulated to sterically impede Y phosphorylation and/or access of SH2 domain proteins to phosphorylated Y180. Although this has not been formally proven, no doubly phosphorylated peptides have been reported in the Phosphosite database [47], suggesting mutually exclusive rather than cooperative phosphorylation at these sites. Cnidaria lack T178 but commonly possess an acidic stretch C-proximally to the cognate of Y180, also commonly including proximal C-terminal S and T potential sites of phosphorylation, resembling the CK2 consensus site of S181 (Figure S2). We propose that a novel functional antecedent to the PGRMC1 Y180 motif evolved in the common cnidarian/bilaterian ancestor.

K193 at the PGRMC1 C-terminus is a consensus SUMOylation site [4], which predates divergence of cnidarians and bilaterians (Figure S2). SUMOylated proteins are frequently nuclear [50], perhaps hinting that the occasional nuclear localisation of PGRMC1 reflects an evolutionarily novel function acquired by early-diverging animals that is conserved in bilateral animals.

### The PGRMC1 Y139 and Y180 combination has been conserved since Cnidaria

Although synapses are present in both Cnidaria and Ctenophora [51], the groups are thought to have independently evolved neurons. Cnidarian and Bilaterian synapses are thought to have evolved from a common ancestor [49]. In our study, Cnidarians were the earliest-diverging animals to acquire the combination of the cognates of PGRMC1 Y139 and Y180, and these are strongly conserved across the Bilateria indicating synapomorphic evolutionary appearance in the common ancestor of those groups. This combination corresponds closely to the presence of ubiquitinated K96 and K137 and consensus SUMOylation site K193 mentioned above. Sabbir has recently demonstrated the inducible presence of PGRMC1 phosphorylation as well as SUMOylation and ubiquitination that affected PGRMC1 stability as well as Tcf/LEF transcriptional activation [52]. By employing sequences of earlier-branching animals (sponges, ctenophores and placozoans) and 88 cnidarian PGRMC sequences we can conclude with high certainty that Y180 arose in the Cnidaria/Bilateria common ancestor not shared with earlier-diverging animals. The position of Y139 was commonly a W in many unicellular holozoans, with occasional examples of substitution for Y in Choanozoa, Ctenophora and Placozoa, however the Y139/Y180 combination provided new animal-specific functionality to PGRMC1 in Cnidaria/Bilateria because it is strongly conserved (Figure S2). Y139 is the less strongly conserved of these two residues, suggesting its function is not as critical to animal biology.

### Animal PGRMC MIHIR regions are predicted to form a coiled-coil protein interaction region

In order to explore possible function of the SH2 motif Y139-containing PGRMC1 MIHIR region we performed low stringency BLAST using human PGRMC1 MIHIR sequence as the search string. Standard BLAST parameters returned only PGRMC1 and PGRMC2 with 100% and 63% similarity and BLAST scores of 82 and 53. However, low stringency BLAST also recognised Neudesin as the next top hit with 48% similarity and a score of 28, followed by a long list of proteins including multiple myosins with BLAST scores above 25. The region of best alignment (PGRMC1 133-164) was then BLASTed against animal species, revealing similar myosin-like motifs from sponge, insects and chordates (Figure 5a). This region was found to have partial predicted coiled-coil character in both PGRMC1 and PGRMC2 (Figure 5b). α-Helical coiled coils are protein-folding and -interaction motifs in which two or more α-helical chains interact to form bundles, typically involving amino acids that exhibit heptad repeats to align on the protein-interactions side of each α-helix. As such, they are associated with protein-protein interactions [53, 54]. The corresponding motif of Myosin 10 was in the coiled-coil region of the protein (Figure 5c). For both PGRMC and myosin motifs, predicted coiled-coil probability was higher in the N-terminal portion of the motif, and reduced in the C-terminal residues.

**Figure 5.**
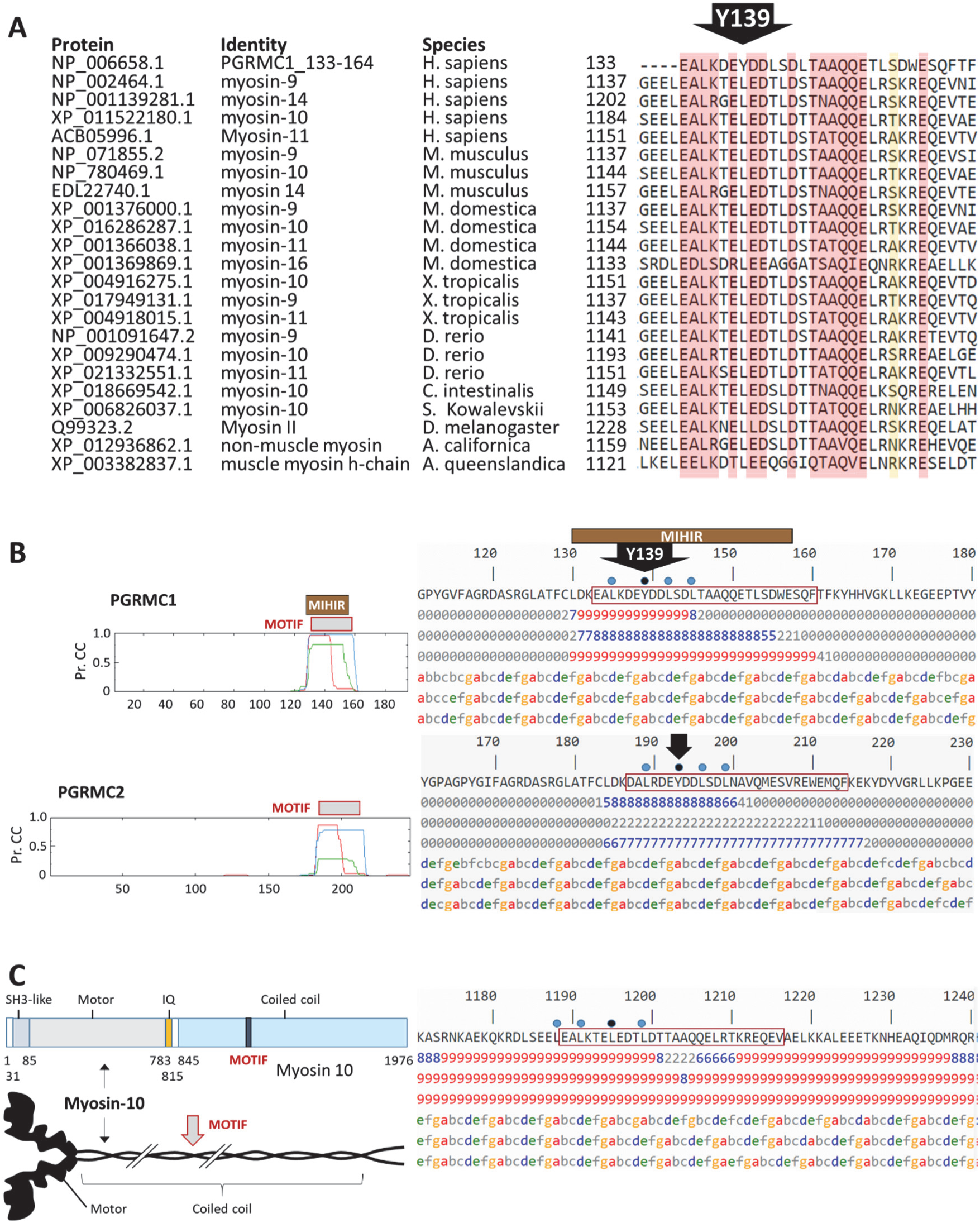
The PGRMC1 MIHIR has predicted coiled-coil character shared with some myosins. **(A)** Alignment of PGRMC1 MIHIR residues 133-164 with selected myosin proteins detected by low stringency BLAST. **(B)** The PGRMC1 and PGRMC2 MIHIR regions contain predicted high propensity to form coiled coil. The images to the left depict the probability for a particular residue to form coiled-coil based upon calculation for surrounding windows of 14 (red), 21 (blue) and 28 (green) residues, generated by the PRABI server. Panels to the right present the numerical depiction of the same result. Numbers under the sequence are the probabilities for coiled-coil formation abbreviated to first digit for windows of 14, 21 and 28 residues (i.e. 9 represents p≥0.9). Letters a-g represent the corresponding coiled coil heptad register. The positons of predicted heptad hydrophobic coiled-coil core residues including PGRMC1 Y139 are indicated. **(C)** The motif from A is present in the coiled-coil region of human Myosin 10. The left side shows the position of the motif in the primary and tertiary structure of the protein. The right side format follows the conventions of B.

We analyzed the probability for coiled-coil formation of this MIHIR motif among selected MAPR proteins by summing the single digit coiled-coil probability scores for each residue in the homologous motif (Figure S3). The motif showed higher propensity for coiled-coil in PGRMC and NENF, with neglibile levels in NEUFC species.

However, coiled-coil probability was not a consistently conserved feature of PGRMC or NENF proteins across species (Figure S3). These data are suggestive of protein-protein interactions occurring through the MIHIR region, possibly via coiled coil interactions in vertebrate PGRMC proteins. Lack of coiled-coil formation does not argue against mediation of protein interactions by the MIHIR. There is no requirement for coiled-coil formation to enable functional protein interactions, as long as the respective interaction surfaces co-evolved compatibly in any given species.

### NENF family

#### N-terminus

The NENF MAPR domain is quite proximal to its transmembrane domain/signal peptide, with no evident systematic patterns observed from early- to late- diverging opisthokonts between transmembrane helix and MAPR domain (Supplemental Information File 2).

#### MAPR domain

In our phylogenetic reconstruction, Holomycota plus Ichthyosporea formed a well-supported clade separate from the remaining Holozoa. Therefore we have denoted all sequences from this clade as “NENF-like”. The major differences between NENF-like and NENF MAPR proteins are color-coded in Figure S4. With regards to accepted rooted phylogeny of the groups concerned (Figure 1A), NENF-like proteins appear to represent an ancestral/pleisiomorphic state for the NENF supergroup. Therefore, the animal-like NENF MAPR domain status represents a synapomorphy that appeared after the common ancestor of icthyosporeans and metazoans (absent from Icthyosporea, present in Choanozoa, Figure S4) from where it was inherited by animals.

#### C-terminus

Cnidaria/Bilateria NENF proteins have acquired a C-terminal extension relative to Porifera, Choanozoa, and early-branching metazoans (Figure 3, Figure S4). The ProteinPredict server predicts a protein interaction region between residues 145-150 (Figure S5). There is no sequence similaritiy between the C-terminal extensions of PGRMC and NENF proteins, indicating probable pronounced functional divergence and specialization of these proteins during early metazoan evolution.

### NEUFC family

#### MAPR domain

There are several conspicuous developments during the evolution of NEUFC. Ctenophora, Cnidaria and bilateria acquired a common histidine at H72 (human NEUFC numbering) in the vicinity of the heme binding pocket, suggesting altered ligand-binding. Animals but not single-celled Holozoa have a greater probability of aspartate at D103, and relative to choanozoa, animals possess a two-residue extension at the C-terminus of the MAPR domain including highly conserved G135 (Figure S6). This region corresponds to a site of predicted protein intetraction (Figure S7).

#### C-terminus

Like all MAPR families, the NEUFC family acquired a C-terminal extension during opisthokont evolution. Unlike PGRMC and NENF families, the NEUFC C-terminus is elongated already in choanoflagellates (Figure 3), relative to earlier-branching holozoans and also fungi. Elements of the evolutionarily newly acquired NEUFC C-terminus are strongly conserved between choanoflagellates and all animals surveyed (Figure S8), implying that this region plays a necessary role in the organismal biology of later-diverging holozoans. It is the largest conserved C-terminus of the animal MAPR family.

Various sites of protein interaction were predicted in the C-terminus by ProteinPredict, as well as a predicted solvent-exposed helix from approximately residues 150-170 (Figure S7A-B). That helical region exhibited a high probability of coiled-coil formation in some but not all NEUFC species sampled (Figure S7C-E). The overall conservation of those residues seems to be more associated with charged residues rather than heptad hydrophobic residues required for coiled-coils. Choanozoans exhibit what appears to be a classical evolutionary intermediary stage between the lower holozoan and fungal state on the one hand, and that of animals on the other (Figure S8). In summary it is highly likely that the NEUFC C-terminus is involved in protein interactions through solvent-accessible residues, however further studies will be required to determine the nature of such interactions, and shed light on the function of NEUFC.

## DISCUSSION

A major finding of this study is that all three animal MAPR families had already diverged in the last common ancestor of the opisthokonts. We detected NENF genes in Choanozoa, unlike Ren at al. [3], demonstrating that all three gene families were present in the common ancestor of Choanozoa and animals. Indeed, Ren et al. concluded that the sequence repertoire for animal MAPR genes likely arose in an ancestral animal sequence, which our results refute. While unicellular opisthokonts with all three genes are rare, one such lineage must have proliferated from the ancestral Opisthokont to give rise to choanoflagellates and animals. Loss of genes in particular lineages is a common observance in opisthokont evolution. For instance, although a common ancestor of the divergent yeasts *S. cerevisiae* and *S. pombe* must have possessed PGRMC, NENF and NEUFC family genes, because the common ancestor of all fungi must have (all three MAPR sub-families are found in the Holomycota (Figure 1B)), both extant organisms possess only a single PGRMC-related Dap1 gene (this study).

Strikingly, the extension of the C-termini of all three MAPR families appeared either before the choanoflagellate divergence or at some point in early animal evolution. For NEUFC the origins of this extension occurred prior to the emergence of the choanoflagellates, the sister group to animals. The PGRMC C-terminus gained an extension before the divergence of Placozoa and Cnidaria, while for NENF this occurred with the Cnidaria/Bilateria common ancestor. There is no sequence similarity between these C-terminal extensions, so this phenomenon represents a further functional divergence between the three MAPR proteins of early animals, associated with the transition to multicellularity and increased organismal complexity. Because little is known about the functions of MAPR proteins, or functional differences between family members [7], cellular functions cannot be ascribed to these novel animal-specific MAPR regions.

### PGRMC1/2 tyrosine phosphorylation motifs

Another major finding of this study is that the combination of SH2 motif phosphoacceptor residues Y139 and Y180 first appeared in the common ancestor of Cnidaria and Bilateria, being absent from Porifera, Ctenophora and Placozoa (Figure S2). That ancestor was among the first animals to possess neurons with Bilaterian-like synapses [49]. PGRMC1 is Y-phosphorylated in synapses [35], and affects synaptic function [14]. The common Cnidaria/Bilateria ancestor, which was probably bilaterally symmetrical, evolved an organizer capable of inducing differentiation of surrounding cells to define embryological body axes and tissue identities that involved gastrulation at an animal pole. That set in motion an orchestrated set of events involving the induced expression of conserved transcription factors such as *brachyury*, *goosecoid* and *foxa* [55]. This is reflected by organizers recognized, for instance, in Cnidaria [56, 57], and Bilateria including Planarian flat worms [58], arthropods [59], spiralian protostomes [60], and of course the deuterostome/chordate Spemann-Mangold organizer [61, 62].

All of the Bilateria *sensu stricto* posses the Y180 motif with adjacent T178 and S181 (Figure S2), all of which residues can be phosphorylated in mammals [25]. Because of the evolutionary appearance of this motif at the same time as the rules governing vertebrate embryological cell-type and tissue differentiation became established, we predict that inappropriate alterations in the phosphorylation status of Y180 in PGRMC1 (or PGRMC2) could impose profound effects on human cells, and therefore could be of potentially monumental clinical importance. Tyrosine phosphorylation is typically caused by induced signal transduction pathways. The signal systems surrounding the regulation of Y180 are likely to feature prominently in human disease.

Cnidarians such as *Hydra* possess a Wnt-dependent head organizer which drives axis specification through a protein gradient [63–65]. The Spemann-Mangold organizer also specifies vertebrate dorso-ventral axis via Wnt-signaling [66]. PGRMC1 is involved in the maintenance of human embryonic stem cell pluripotency via regulation of the Wnt pathway [28]. Based upon the strongly conserved coincident presence of the Y139 and Y180 motifs shared between Bilateria and Cnidaria we hypothesize that PGRMC1 phosphorylation might also play an important role in the cnidarian organizer. Furthermore, PGRMC1 may ancestrally be involved in the transition from the single morphotype protist state to the development of the collective of multiple states of differentiated morphology that is characteristic of the clonal metazoan condition, where the ability to phosphorylate Y139/Y180 could have been associated with the evolutionary origin of organizer function and tissue differentiation. PGRMC1 is known to be SUMOYlated which affects TCF/LEF-driven transcription [52, 67]. TCF/LEF is probably the exclusive conduit for Wnt signalling [68]. Taken together, PGRMC1 is strongly implicated in the evolution of animal organizer function with profound potential to influence animal cell differentiation status and its plasticity (e.g. cancer) [7, 52, 69].

In accompanying papers we show that mutational manipulation of the phosphorylation status of the Y180 motif dramatically affects cultured cancer cell morphology, PI3K/Akt signaling activity, mitochondrial form and function, glycolytic/energy metabolism, and ability to form tumours in MIA PaCa-2 pancreatic cancer cells. This is reflected in altered cell metabolites, genomic stability, and dramatic changes in genomic CpG methylation [70, 71], which are all consistent with PGRMC1 invovelment in organizer biology.

### PGRMC and NENF involvement in neurology

In addition to the combination of PGRMC1 Y139/Y180 discussed above, the C-terminal extension of NENF was also acquired in the Cnidaria/Bilateria common ancestor, coincident with the appearance of neurons with synapses. NENF neurotrophism [16] and PGRMC1 axonal guidance [8, 9] are at least superficially similar functions.

For reasons such as these we were expecting to find that animals inherited a single MAPR gene from their unicellular ancestors, which diverged into three families during metazoan evolution, with e.g. ancestral neural-related functions having undergone functional specialization following gene duplication. While this manuscript was in preparation Ren at al. [3] concluded that choanoflagellates and the first metazoans contained only PGRMC and NEUFC genes. The findings that all three MAPR families were already distinct in the ancestral opisthokont strongly negates both hypotheses. However, the observation remains that PGRMC1 and NENF both acquired new conserved functional features at the emergence of Cnidaria/Bilateria common ancestor, correlating with the evolution of nerve synapses, despite having been inherited into animals as separate genes from a unicellular ancestor. There seems to have been some feature of MAPR biology which was important in the ancestors of animals and which favoured the subsequent evolution of a cnidarian/bilaterian grade of body organization that included synapses. We hypothesize that at least PGRMC1 Y180 was crucial in the latter.

In this light, it is interesting that in NENF knockout mice Novais et al. [72] observed decreased hippocampal levels of the steroids progesterone and allopregnenolone (APα), and increased levels of 3-oxo-5-alpha-steroid 4-dehydrogenases 2 and 3 (which convert progesterone to APα) and gamma-aminobutyric acid A (GABA) receptor delta (GABRD), an important neuromodulator which is positively allosterically regulated by APα [73]. From yeast (phylogenetically disparate *S. pombe* and *S. cerevisiae*) to mammals PGRMC1-like proteins interact with lanosterol-14-demethylase (CYP51A1) [5, 36, 74], the first enzyme in sterol metabolism leading to animal cholesterol. This situation is consistent with a model where PGRMC proteins in unicellular opisthokonts may have been involved in sterol synthesis pathways for the production of 3-ketosterones such as progesterone, whereas NENF may have regulated the conversion of 3-ketosterones to other biologically active steroids, both of which were required for animals to evolve. i.e. where a PGRMC/NENF gene divergence led to new steroid-associated functions, albeit that we are unaware of 3-keto-steroids being attested in earlier diverging opisthokonts. This hypothesis would unite both these MAPR protein functions with an ancestral role in steroid biology. Consistently with a possible ancestral steroidogenic role of MAPR proteins, NEUFC attenuation led to reduced levels of CYP51A1 in HeLa cells, the same enzyme with which PGRMC1 interacts [5, 75], reinforcing a proposed ancestral association between MAPR proteins and sterol biology. The hypothesis predicts that MAPR gene divergence in the opisthokont lineage leading to animals involved functional diversification towards new pathways of sterol production, among other outcomes including the multiple attested functions of PGRMC1 (reviewed elsewhere [7]).

### Is the MIHIR a protein-interaction motif?

We identified a motif with probable coiled-coil characteristics in the MIHIR sequence of both PGRMC1 and PGRMC2, as well as similar sequences in some myosins. The motif was predicted with high probability to form a short coiled-coil at its N-terminus, and lower probability at its C-terminus. Of particular interest was the observation that Y139 formed part of the hydrophobic heptad repeat required for interaction of adjacent helices in the coiled-coil. In lower Holozoans Y139 was commonly a tryptophan, another large bulky hydrophobic residue. The putative involvement of Y139 in a coiled-coil interaction presents immediate connotations when the residue is phosphorylated. Not only would it be unable to interact with coiled-coil interaction partners, but it could then interact with a new set of SH2 domain-containing proteins in the tyrosine-phosphorylated form. This suggests the acquisition of a regulatory switch in MIHIR functionality by PGRMC genes in the Cnidaria/Bilateria common ancestor.

It should be noted that this region of PGRMC1 in the 4X8Y crystal structure exhibited a high B or scatter factor, indicating relatively poor mapping of electron density to amino acid sequence [76], as indicative of a region of low structural stability in the crystal. We therefore conclude that the MIHIR region should be able to rapidly sample many conformations in solution, which is compatible with availability to form protein-protein interactions.

In myosins, the analogous motif is found within the rod-like coiled-coil region, and represents an area where the probability of coiled coil interaction is diminished, as shown for myosin 10 in Figure 5b. Such coiled-coil regions with weaker stability, often including disruption of the requisite heptad repeat coiled-coil motif, have been proposed to form sites of binding to potential target interacting proteins [77]. A phylogenetic survey of some MAPR proteins from opisthokonts (Figure 5f-g) showed that high predicted probability of coiled-coil formation was conserved in animals (except the sea anemone *Aiptasia pallida*), as well as in *Monosiga brevicollis* from the choanoflagellate sister group of animals. The degree of coiled-coil propensity was weaker in earlier-branching single celled opisthokonts, being minimaly present in the yeast *S. pombe* (Figure 5f-g). We predict that the MAPR MIHIR sequence, a highly conserved early eukaryotic innovation, enables protein interactions not shared with other cytb_5_ proteins, which may or may not involve coiled-coil formation. Association with the actin/myosin cytoskeleton through the MIHIR motif may be related to the membrane-trafficking functions of PGRMC1.

### A PGRMC1 RGD integrin-interaction motif?

We observe a potential vertebrate RGD motif in what is conventionally considered the cytoplasmic region of PGRMC1 of some vertebrates, being absent in the turtles, birds and marsupials studied (Figure 4). This assignment is tenuous because RGD motifs are predicted to be involved with extracellular integrin interactions [78]. Integrins are important in many processes, including the regulation of synaptic plasticity and memory [79]. The RGD motif would need to be extracellular to be functional. However, the conventionally cytoplasmic C-terminal region of PGRMC1 is extracellular in several attested situations, including synapses, where PGRMC1 is involved in a mechanism that affects synaptic plasticity [14], in pluripotent stem cells [80], where PGRMC1 is associated with the maintenance of pluripotency involving the Wnt/beta-catenin pathway [28], and in cancer cells, where PGRMC1 may be secreted by the exosome pathway [81].

PGRMC1 is documented as an exosomal protein in the Exocarta database (http://www.exocarta.org/) [82], however it remains unclear whether it may be translated with an alternative membrane topology as a transmembrane protein [7, 83]. We surmise that the likelihood of a conserved functional RGD integrin-interaction motif in vertebrates would be much greater if the protein exhibits alternative translational topology as a transmembrane protein, rather than as a secreted protein. In view of the association of PGRMC1 with fertility [34, 84, 85], it is conveivable that this motif is involved in the evolution of post-fertilization vertebrate embryology, perhaps involving the relationship between amniotic sac and eggshell, or other major features related to differences in vertebrate embryology and oocyte/egg biology between these groups.

Positively charged residues cytoplasmically proximal to transmembrane signal peptide helices are critical in determining the orientation of transmembrane proteins [86]. There is potential to alter membrane topology of PGRMC1 by post-translational modification of K44 or R47 during translation. It is known that R47 can be methylated (Phosphosite action?id=5744). This may be relevant to the observation of extracellular PGRMC1 C-terminus in the synaptic extracellular space of neurons [14], potentially via regulated non-conventional orientiation during translation. However we stress that in the absence of validated integrin interaction data, the presence of a functional RGD motif in vertebrate PGRMC1 must be considered cautiously.

### Conclusions

We initiated this study to try to provide a systematic platform to understand the reported multifunctionality of PGRMC1. Surprisingly, we discovered that members of all three animal MAPR families had originated prior to the origin of opisthokonts. Although many protist species have lost one or more MAPR genes, the common ancestor of each major opisthokont branch leading to animals must have possessed all three MAPR families, evidencing a degree of conservation which indicates that each family performs separate unicellular functions that were essential in the evolution of the holozoan and holomycotan subfamilies of Opisthokonts. We are unaware of a major eukaryotic group which does not possess at least one MAPR gene, or is not thought to have ancestrally done so. Some parasitic groups are thought to have secondarily lost MAPR genes [83].

We propose that the attested PGRMC1 membrane-trafficking function may be related to the eukaryotic acquisition of the MAPR-defining MIHIR region. In a separate study (submitted) we show that the MIHIR is a eukaryotic development, and propose that the ancestral MAPR protein was one of the pivotal proteins involved in the development of the first truly eukaryotic cell, with roles in the evolution of steroid biology.

Mitochondria require cholesterol to increase membrane potential across the inner membrane, and many of PGRMC1’s functions involve steroid biology [83]. Interaction of the eukaryotic MIHIR with the actin cytoskeleton may have been involved in the origin of membrane trafficking that was necessary to target cytoplasmically-synthesized steroids to the proto-mitochondrion before the endosymbiont could lose genes to the nucleus. Additional or parallel to such ancient roles, we hypothezise that the early evolutionary diaspora of eukaryotic diversity involved functional specialisation of at least three opisthokont MAPR genes. The ancestor of Opisthokonta already contained the tripartite MAPR repertoire, although multiple unicellular species have either variously lost MAPR genes or we were not able to find them. The transition from pre-choanozoan to animal multicellularity involved enlargement of the C-terminus in all three MAPR families, commencing with the common ancestors of Bilateria and Placozoa (PGRMC), Cnidaria (NENF), and Choanozoa (NEUFC), respectively (Figure 3). The PGRMC gene acquired the combination of Y139 and Y180 at the stage of evolution of the common ancestor of Cnidaria and Bilateria, concommittantly with the appearance of synapsed nerves and prior to the evolution of Bilateria *sensu stricto* and the associated new mechanisms for embryological body pattern formation and tissue differentiation. Our study suggests a stratified acquisition of PGRMC1 functions, and points towards potentially dramatic effects of cell differentiation status if this ancient axis of PGRMC1 tyrosine phosphorylation is perturbed in disease processes. Results presented in accompanying papers strongly support this hypothesis [70, 71]. These combined findings signpost the direction for productive future studies, which was our aim.

## MATERIALS AND METHODS

### Identification of MAPR proteins

The human sequences of PGRMC1, NENF and NEUFC were used in an initial BLASTP search against the NCBI non-redundant protein sequence database (using default NCBI BLASTp settings), retaining the best hit per organism. Additional sequences were identified by using the same queries in BLASTp searches (evalue 1e-25) against a custom database as described [44], and further expanded using newly available unicellular opisthokont datasets (*Parvularia atlantis* and *Chromospahera perkinsii*, available from http://multicellgenome.com) as well as additional fungal datasets (downloaded from https://genome.jgi.doe.gov/mycocosm/home). The dataset was extended for the placozoan *Hoilungia hongkongensis*, three classes of Porifera, Ctenophora, Cnidaria and also Choanoflagellata by BLAST searches against a set of non-bilaterian proteomes that have been previously established [87, 88]. BLASTP searches were performed with specifying an evalue of 1e-10 and otherwise default settings using the human NENF, NEUFC and PGRMC1 as well as the initially identified placozoan *Trichoplax adhaerens* [89] NEUFC and PGRMC protein sequences. Specimens of the calcareous sponge *Pericharax orientalis* were collected from Dunk Island Mission beach in 2016 (under the authorization CMES59 provided by James Cook University). MAPR sequences of *Pericharax* were retrieved from a draft assembled transcriptome (Adamski et al. in prep). BLASTP searches using mammalian sequences as query allowed the identification of the *P. orientalis* homologs. The MAPR identity of all proteins used was verified by sequence alignment and confirmation of the presence of a MIHIR.

### Phylogenetic reconstruction

All MAPR sequences described above were initially aligned with MAFFT v. 7.212 L-INS-i [90]. Ambiguously aligned positions were trimmed off with trimAL v. 1.2 [91] using a gap threshold of 20% and a tree was calculated using FastTree v. 2.1.7 with default options [92]. The resulting phylogeny and the underlying alignment were manually inspected and obvious contaminations, duplicates and paralogs were removed. The cleaned, unaligned sequences were then subjected to filtering with PREQUAL [93] using the default options to remove non-homologous residues introduced by poor-quality sequences, followed by alignment with MAFFT G-INS-i using the VSM option (--unalignlevel 0.6) [94] to control over-alignment. The alignments were subjected to Divvier [95] using the -divvygap option to improve homology inference before removing ambiguously aligned sites with trimAl v. 1.2 (-gt 0.01). We then extracted the region from position 46 to 139 (94 amino acid residues (aa), relative to human PGRMC1), that is conserved in all taxa of our alignment, from the trimmed alignment and performed the tree reconstruction based on this central conserved region only. Final trees were calculated with IQ-TREE v. 1.6.5 [96], using the -mset option to restrict model selection (to DAYHOFF, DCMUT, JTT, WAG, VT, BLOSUM62, LG, PMB, JTTDCMUT; model selected: LG+R5) for ModelFinder [97], while branch support was assessed with 1000 ultrafast bootstrap replicates [98]. We also prepared a conservative phylogeny without performing PREQUAL filtering prior to sequence alignment or Divvier analysis after alignment, followed by stringent trimming with trimAl v. 1.2 (-gt 0.8), resulting in an alignment of 126 aa length. The tree was calculated using IQ-TREE and ModelFinder as described above (model selected: LG+R7), branch support was assessed with 1000 ultrafast bootstrap replicates.

With both approaches, we observed a similar topology, particularly the conspicuous split within the NENF clade: while cnidaria, sponges and choanoflagellates formed a well-supported holozoan clade with the metazoan NENF representatives, the holozoan lineage of ichthyosporeans grouped with high support with the holomycota (fungi and nucleariids).

The alignment of selected sequences chosen to provide indicative phylogentic representation of land vertebrates in Figure 4 was made with the Computational Biology Research Consortium (CBRC) MAFFT platform (https://mafft.cbrc.jp/) using the L-INS-i strategy [99].

### Other database platform queries

The same alignment used for tree reconstruction in Figure 1 has been used to generate logo plots to identify conserved regions in groups of interest. The untrimmed 3-protein-alignment was split into three separate alignments for each protein (PGRMC, NENF and NEUFC), followd by removal of gap-only columns in each alignment. For the alignments in Figures S1-S5, a choanoflagellate-specific insert indicated in Figure 2 has been removed. The species corresponding to phylogenetic groups labelled in Figures S2, S4, S6 and S8 were entered to Weblogo separately. Logo plot representation of consensus sequences were generated with the WebLogo platform (http://weblogo.berkeley.edu/) [100], as described in respective figure legends.

Low stringency protein sequence BLAST was performed using human PGRMC1 search string CLDKEALKDEYDDLSDLTAAQQETLSDWESQFTFKYHH with NCBI BLASTp (https://blast.ncbi.nlm.nih.gov), employing word size 3, expect threshold 1000, organism restricted to Homo sapiens (taxid:9606), 500 maximum target sequences, and with all other parameters set to default values. Subsequent BLAST queries targeted the organisms from Figure 5A. Coiled-coil prediction [101] was performed by the PRABI (Pôle Rhône-Alpes de Bioinformatique) server (https://npsa-prabi.ibcp.fr/). Sites of observed phosphorylation were obtained from Phosphosite (www.phosphosite.org) [102] or UniProt (www.uniprot.org). Protein interaction sites were predicted with the PredictProtein server (www.predictprotein.org) [103]. RGD protein interaction motif detection was by the ISIS (interaction sites identified from sequence) [104] function of ProteinPredict.

## LIST OF ABBREVIATIONS

CK2: casein kinase 2
CNS: central nervous system
cyP450: cytochrome P450
cytb_5_: cytochrome b_5_
MAPR: membrane-associated progesterone receptor
MIHIR: membrane-associated progesterone receptor-specific interhelical insert region
NENF: neudesin
NEUFC: Neuferricin
P4: progesterone
PGRMC: progesterone receptor membrane component 2
PGRMC1: progesterone receptor membrane component 1
SH2: Src homology 2
SH3: Src homology 3

## DECLARATIONS

### Ethics approval and consent to participate

Not applicable.

### Consent for publication

Not applicable.

### Availability of data and materials

The raw tree files in newick, colored trees with taxon information in pdf format, and underlying trimmed alignments corresponding to both phylogenetic reconstructions have been deposited to figshare repository doi: 10.6084/m9.figshare.9162164.

### Authors’ contributions

M.A.C. conceived the project, performed WebLogo studies, and wrote the first draft of the manuscript. E.H. performed MAPR sequence identification, sequence alignment and curation and phylogenetic reconstruction with advice from P.J.K. M.E. retrieved additional choanoflagellate, Porifera, Ctenophora, Placozoa and Cnidaria protein sequences. S.A.V.F. sequenced the Pericharax orientalis genome, under supervision from D.J.M. All authors read and provided critical comment to the manuscript.

## Acknowledgements

Not applicable.

## Supplemental Figures

**Figure S1.**
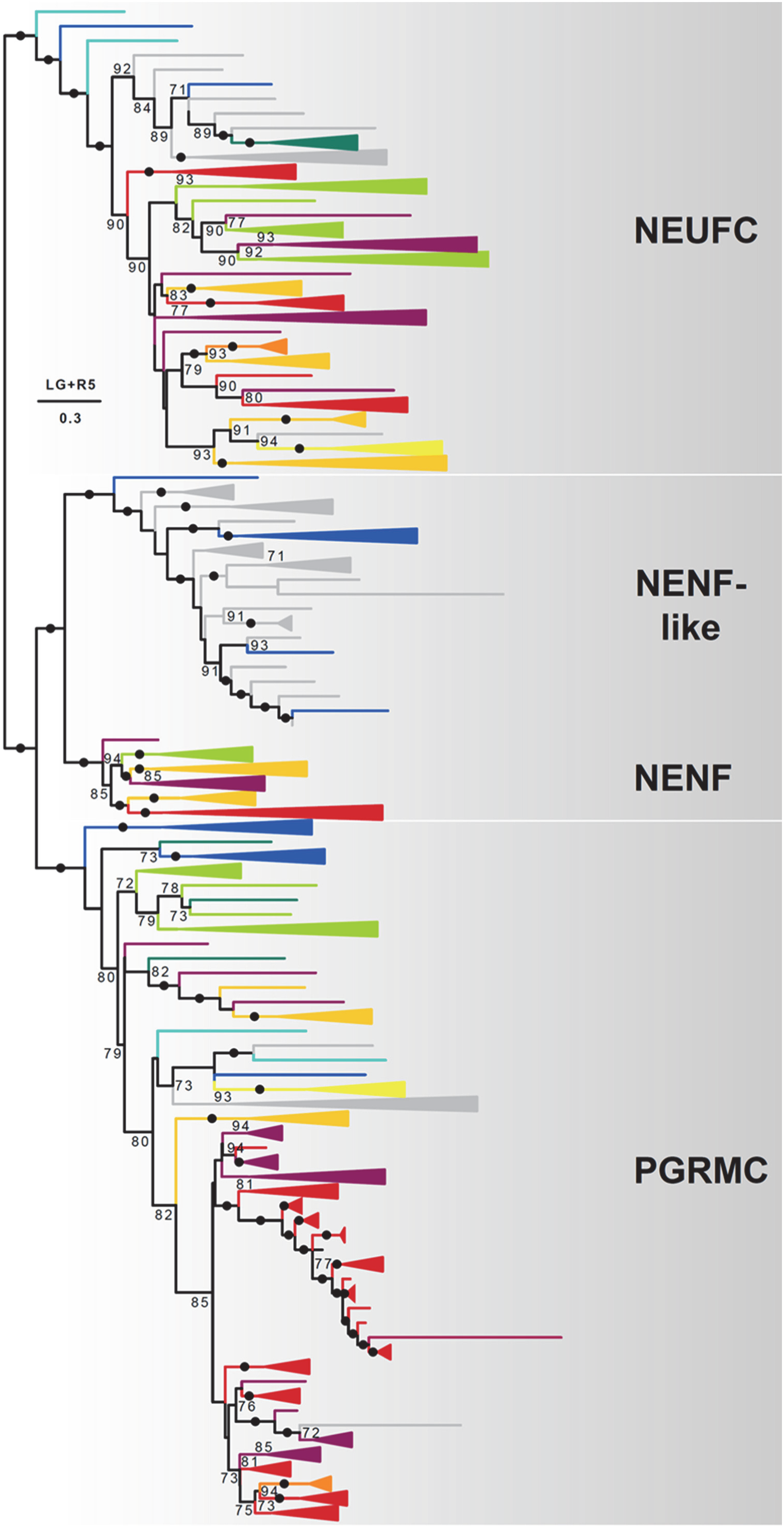
Phylogenetic reconstruction of MAPR proteins in opisthokonts. Phylogeny of 3 types of MAPR proteins in opisthokonts. Opisthokont lineages are indicated by colored branches/collapsed clades and nomenclature following Figure 1B. The scale bar and the number beneath it indicate the estimated number of substitutions per site, above the scale bar the model for tree reconstruction is indicated. Node support was calculated using 1000 ultrafast bootstrap (UFBoot) replicates. Only support values >70% are shown, black dots indicated support values of ≥ 95%.

**Figure S2.**
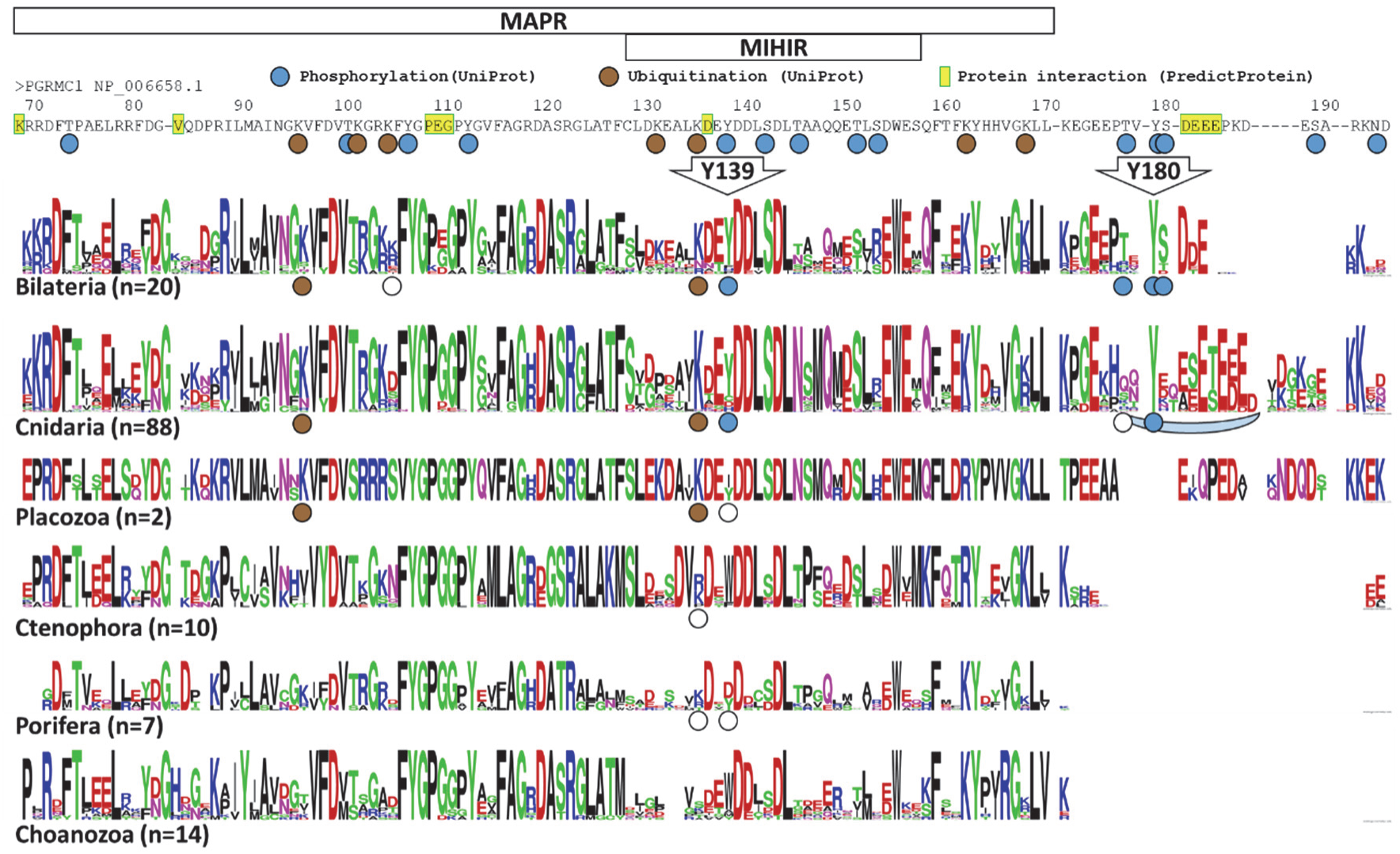
PGRMC MAPR and C-terminus Logo plots. Conventions follow Figure 2. The human PGRMC1 sequence is provided at the top for reference. Logo plots were constructed from the alignment of Supplemental Information File 1.

**Figure S3.**
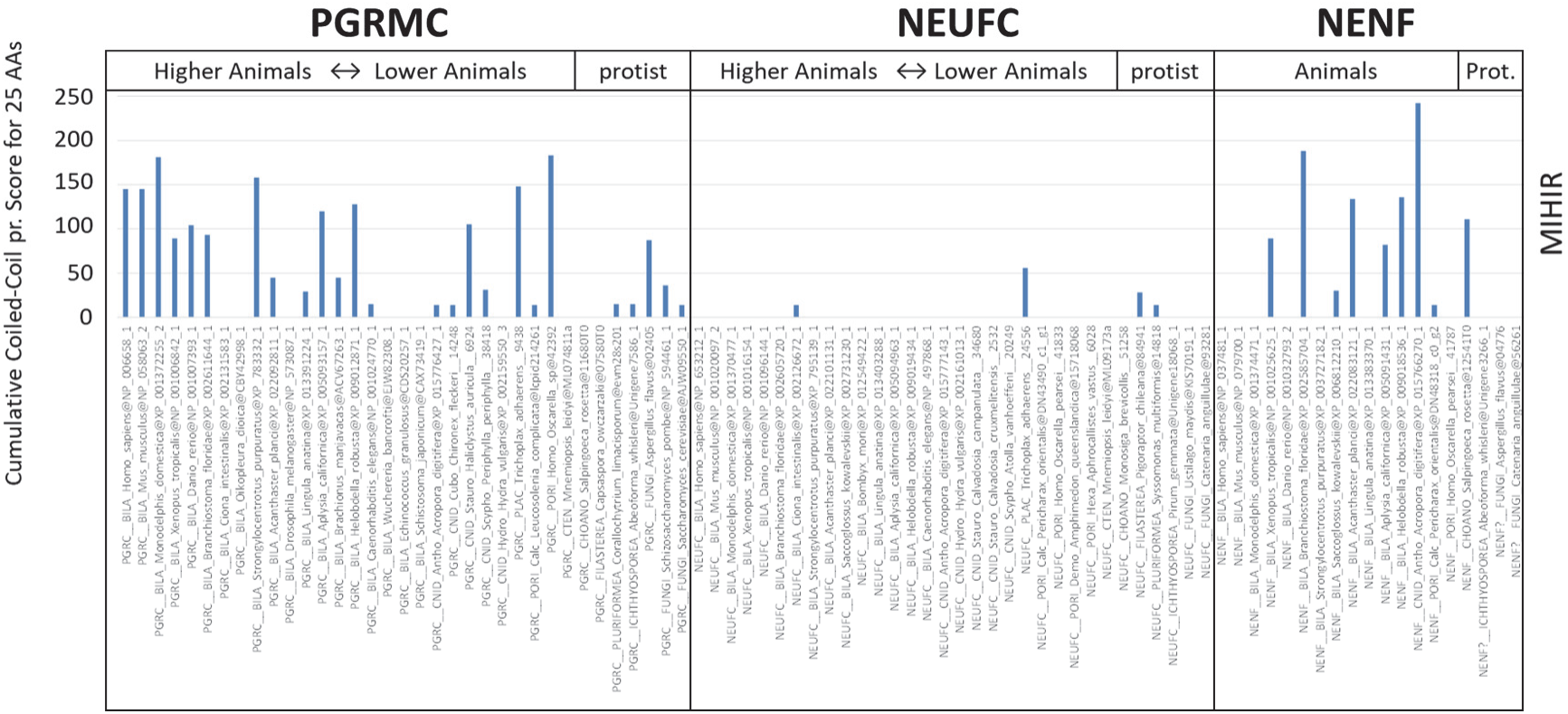
Coiled-coil probability of the MIHIR motif from selected MAPR sequences. The cumulative predicted probability for coiled-coil (Cumulative coiled-coil Pr.) of 20 residues of each sequence aligned with PGRMC1 MIHIR residues TFCLDKEALKDEYDDLSDLT. By way of example, for human PGRMC1 the score was generated by 0+0+2+7+9+9+9+9+9+9+9+9+9+9+9+9+9+9+8+2=145 as calculated by the PRABI server.

**Figure S4.**
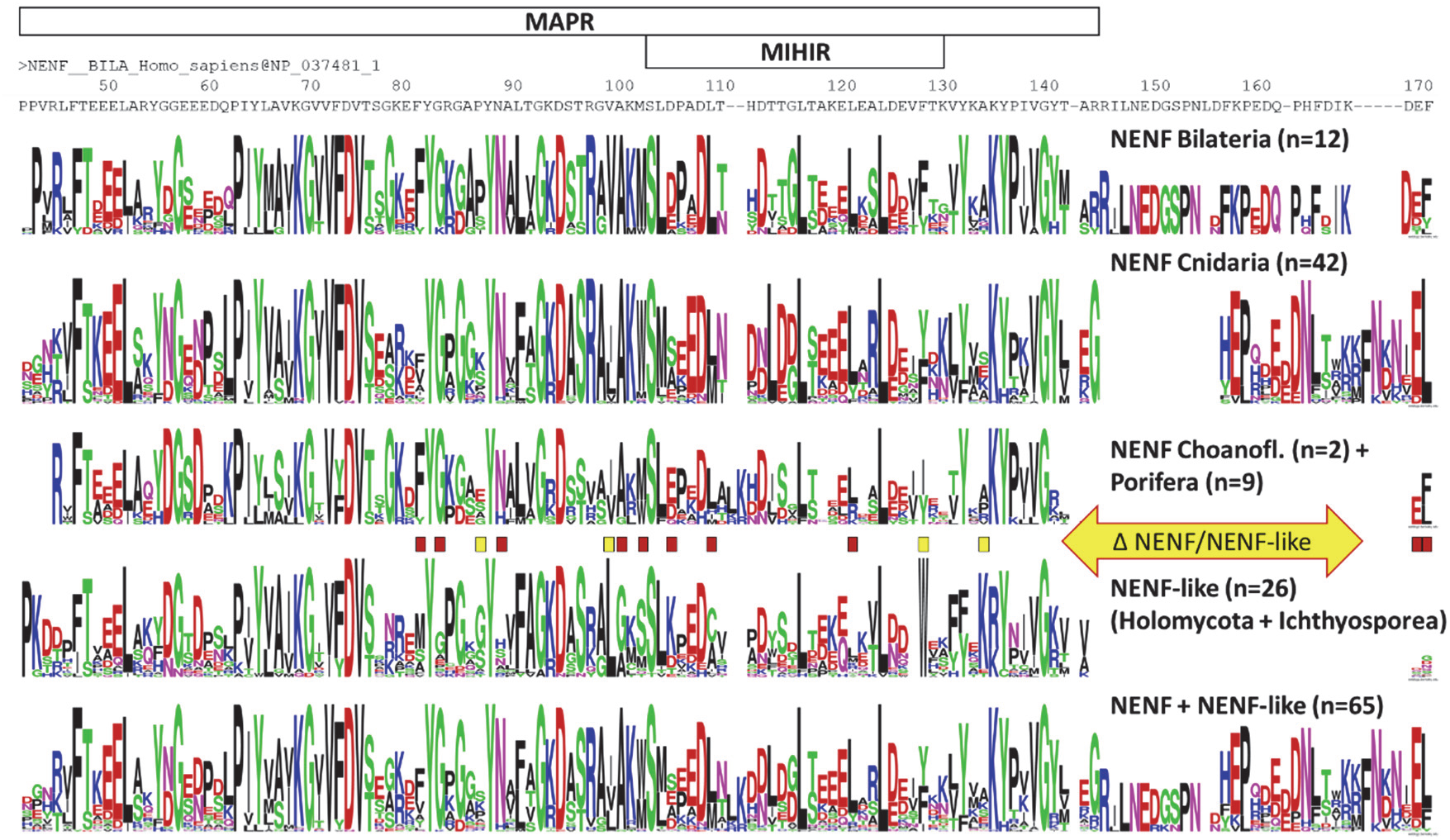
NENF MAPR and C-terminus Logo plot. Conventions follow previous figures. The human NENF sequence is provided at the top for reference. Logo plots were constructed from the alignment of Supplemental Information File 3. The positions of major differences between NENF and NENF-like consenus sequences are indicated (Δ NENF/NENF-like). Red (dark) boxes represent residues where differences are more conserved in NENF, whereas yellow (light) boxes represent residues where differences are more conserved in NENF-like.

**Figure S5.**
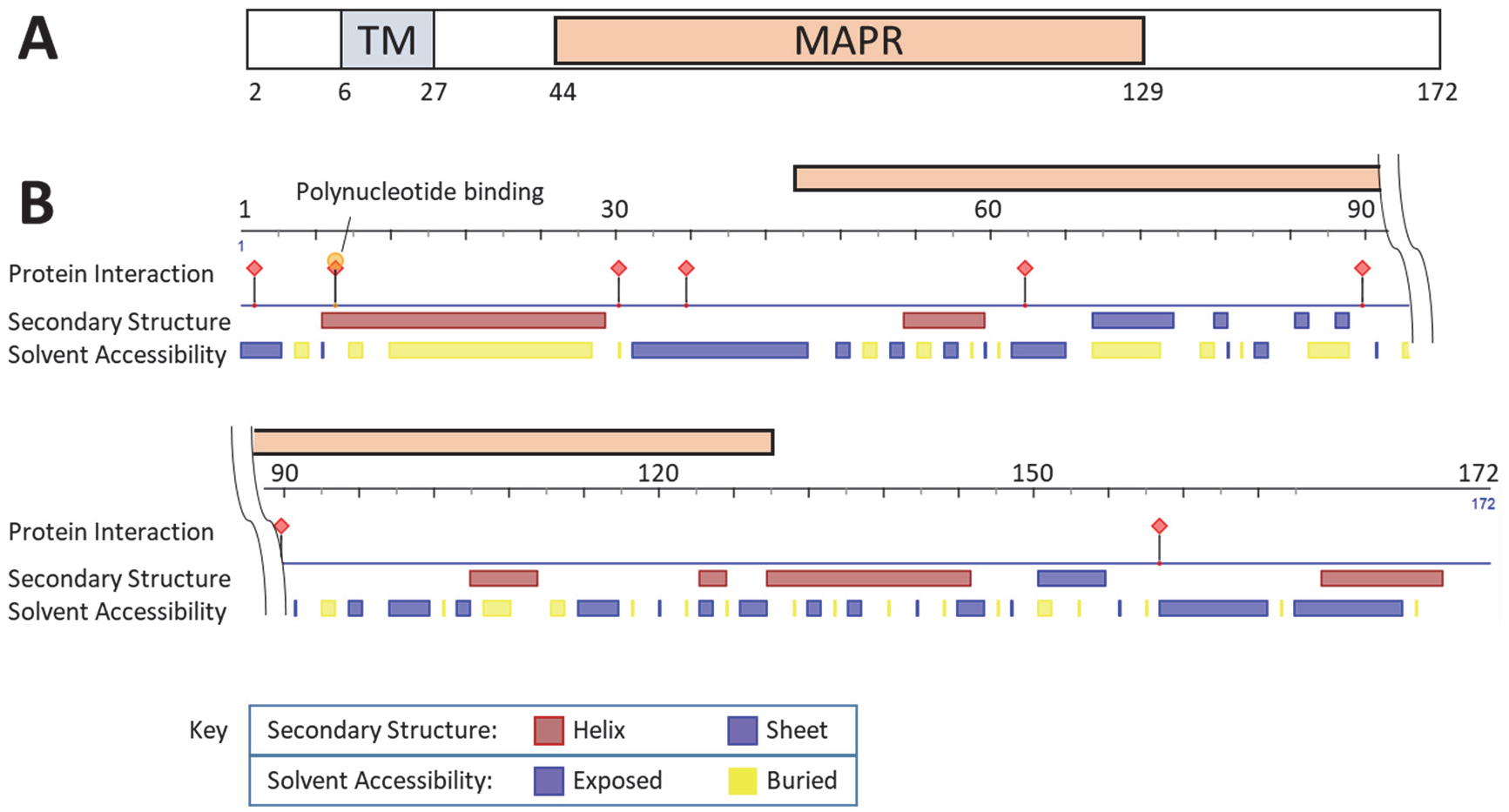
NENF predicted features. A) Schematic representation of Human NENF, showing the position of the transmembrane helix (TM) and MAPR domain B) Graphical depiction produced by the ProteinPredict server, showing sites of predicted protein interactions, secondary structure, and solvent accessibility. A single site of predicted polynucleotide binding is indicated at residue 7.

**Figure S6.**
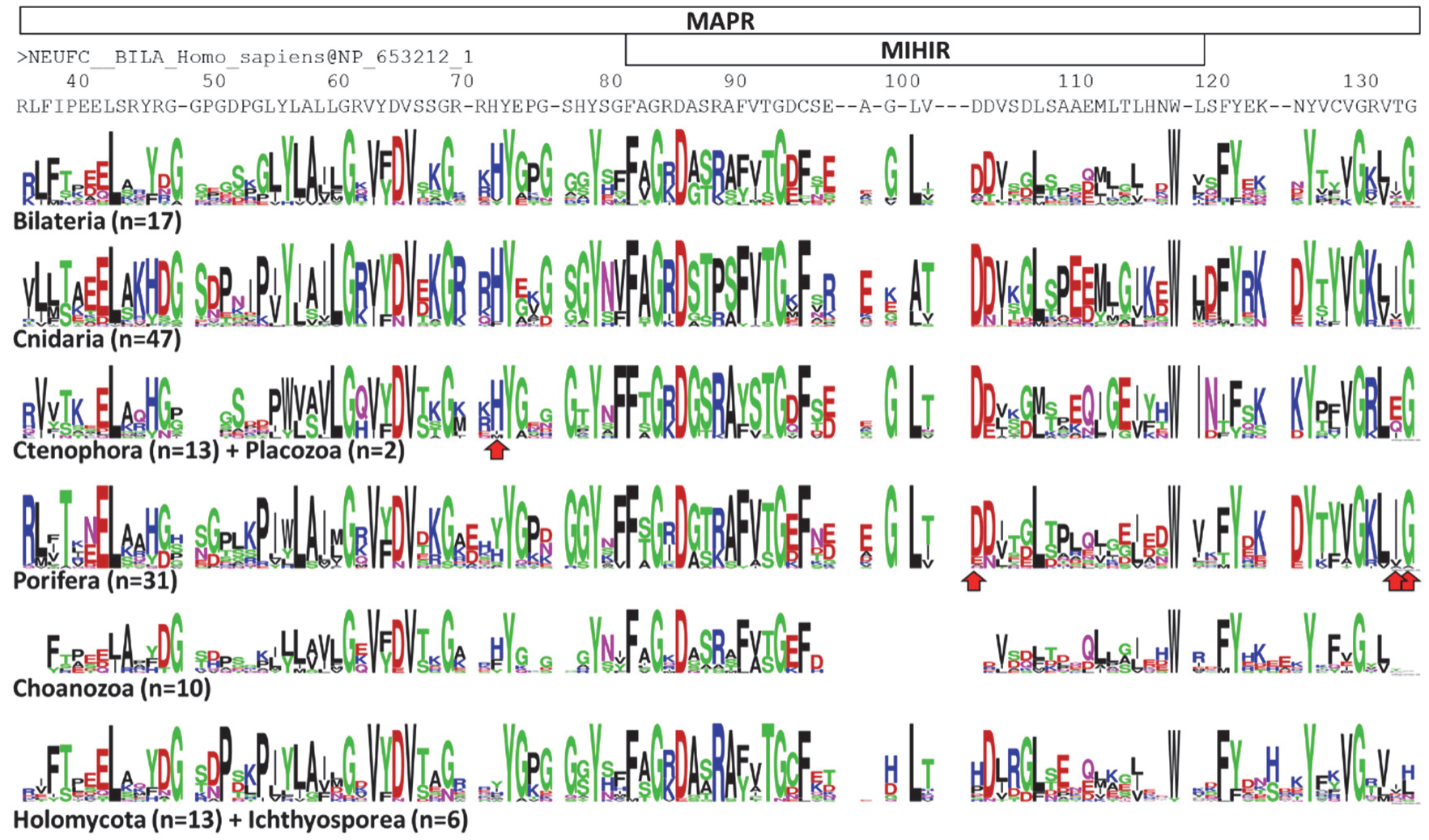
NEUFC MAPR domain Logo plot. Conventions follow previous figures. The human NEUFC sequence is provided at the top for reference. Logo plots were constructed from the alignment of Supplemental Information File 4.

**Figure S7.**
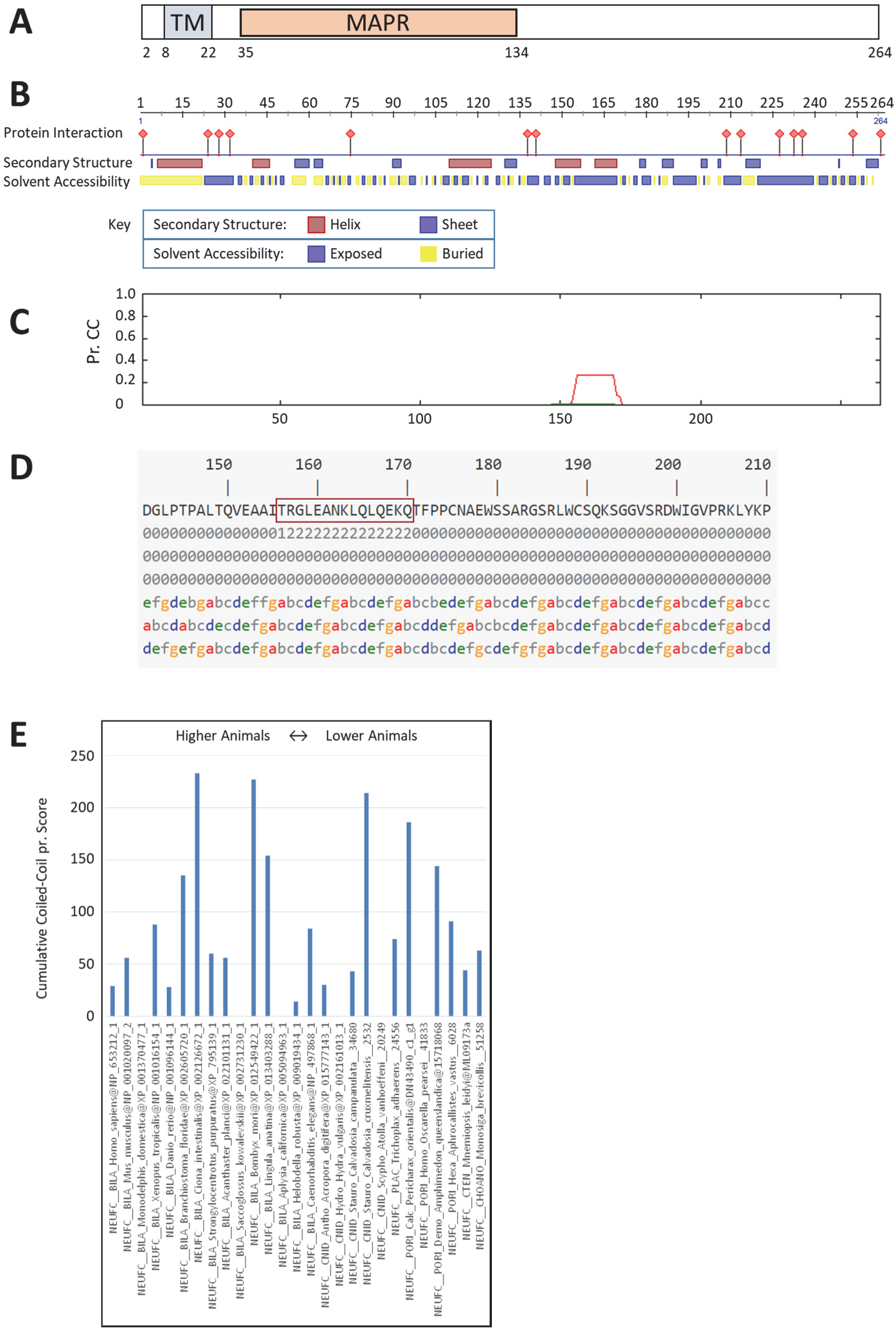
NEUFC predicted features. A) Schematic representation of human NEUFC. Conventions follow Figure S5A. B) Graphical depiction produced by the ProteinPredict server, showing sites of predicted protein interactions, secondary structure, and solvent accessibility. C) Graphical depiction of predicted probability of forming coiled-coil generated by the PRABI server, following Figure 5B. D) Numerical output of the results from C), following Figure 5B, with amino acid sequence boxed. E) Cumulative predicted coiled-coil scores for the residues aligned with human NEUFC TRGLEANKLQLQEKQ from D), following the convention of Figure S3.

**Figure S8.**
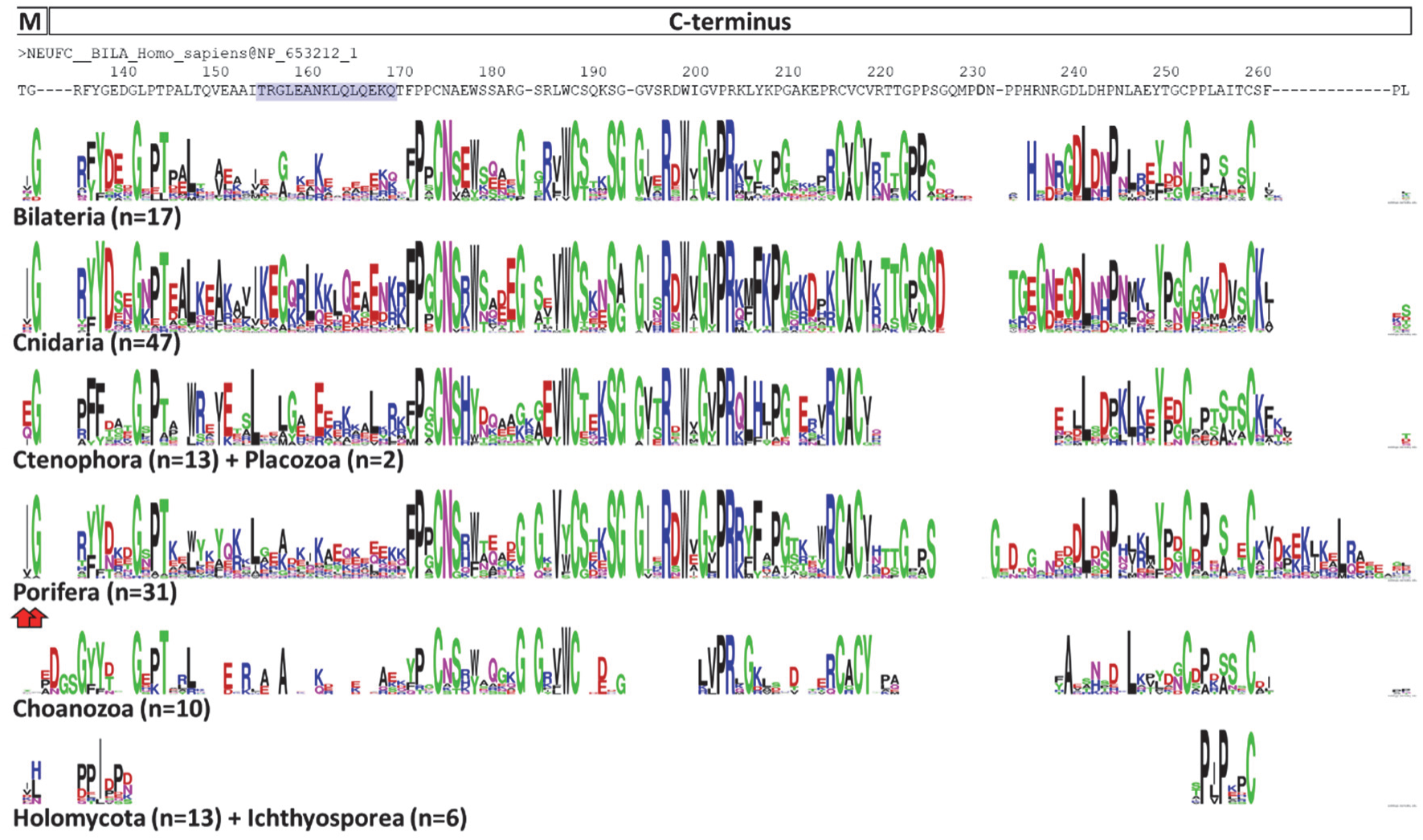
NEUFC C-terminus Logo plots. Conventions follow previous figures. The human NEUFC sequence is provided at the top for reference. Shaded residues (purple) are those referred to in Figure S7C-E. Logo plots were constructed from the alignment of Supplemental Information File 5. Boxes at the top denote two residues fdorm the MAPR domain (M) and the C-terminus of the NEUFC consensus Logo plot (C-terminus).

## Supporting Information files

**Supplemental Information File 1.** PGRMC MAPR domain and C-terminus alignment used for Logo plots of Figure S2. (PGRMC-LOGO_20181026.fasta)

**Supplemental Information File 2.** NENF N-terminal alignment. No Logo plot produced. (NENF-BIG_20181026.fasta)

**Supplemental Information File 3.** NENF MAPR domain and C-terminal alignment used for for Logo plots of Figure S4. (NENF-small_20181026.fasta)

**Supplemental Information File 4.** NEUFC MAPR domain alignment used for for Logo plots of Figure S6. (NEUFC-MAPR_20181026.fasta)

**Supplemental Information File 5.** NEUFC C-terminal alignment used for for Logo plots of Figure S8. (NEUFC-C-term_20181026.fasta)

